# Diminished pre-stimulus alpha-lateralization suggests compromised self-initiated attentional control of auditory processing in old age

**DOI:** 10.1101/439919

**Authors:** Martin J. Dahl, Liesa Ilg, Shu-Chen Li, Susanne Passow, Markus Werkle-Bergner

## Abstract

Older adults experience difficulties in daily situations that require flexible information selection in the presence of multiple competing sensory inputs, like for instance multi-talker situations. Modulations of rhythmic neural activity in the alpha–beta (8–30 Hz) frequency range in posterior brain areas have been established as a cross-modal neural correlate of selective attention. However, research linking compromised auditory selective attention to changes in rhythmic neural activity in aging is sparse.

We tested younger (n = 25; 22–35 years) and older adults (n = 26; 63–76 years) in an attention modulated dichotic listening task. In this, two streams of highly similar auditory input were simultaneously presented to participants′ both ears (i.e., dichotically) while attention had to be focused on the input to only one ear (i.e. target) and the other, distracting information had to be ignored.

We here demonstrate a link between severely compromised auditory selective attention in aging and a partial reorganization of attention-related rhythmic neural responses. In particular, in old age we observed a shift from a self-initiated, preparatory modulation of alpha rhythmic activity to an externally driven, response in the alpha–beta range. Critically, moment-to-moment fluctuations in the age-specific patterns of self-initiated and externally driven lateralized rhythmic activity were tied to selective attention. We conclude that adult age difference in spatial selective attention likely derive from a functional reorganization of rhythmic neural activity within the aging brain.

## Introduction

Daily communication occurs in challenging environments: A multitude of simultaneous auditory signals compete for limited processing resources and selective attention is needed to successfully separate relevant from irrelevant information. Particularly elderly listeners experience difficulties in multi-talker situations, which derive not only from a decline in perceptual abilities but also from impairments in attentional mechanisms (e.g., Anderson, White-Schwoch, Parbery-Clark, & Kraus, 2013; Getzmann, Golob, & Wascher, 2016; Passow et al., 2012, 2014; Westerhausen, Bless, Passow, Kompus, & Hugdahl, 2015). More specifically, older adults often demonstrate an age-graded shift from internally to externally driven attentional control that is associated with diminished performance in situations requiring flexible adaptation to ongoing task demands (Lindenberger & Mayr, 2014), such as in daily social interactions. Given that self-initiated control is demanding (Craik & Bialystok, 2006), increased reliance on external, environmental support may itself reflect a strategic adaptation to diminishing attentional resources with age (Velanova, Lustig, Jacoby, & Buckner, 2006). But while reliance on environmental support might be a beneficial compensatory strategy in many contexts (Lindenberger, Lövdén, Schellenbach, Li, & Kruüger, 2008; Nehmer, Lindenberger, & Steinhagen-Thiessen, 2010), a stronger dependency on external inputs may decrease the ability to flexibly route information selection in the presence of multiple competing sensory inputs, such as in multi-talker situations. In sum, a reduced capacity for internal guidance of attentional selection and an increased reliance on external inputs may specifically impair older adults′ abilities to actively engage in daily social interactions that comprise a multitude of competing sensory signals. However, the neural signatures of reduced internal control of auditory attention in older adults are not yet clearly identified.

Across modalities, attentional engagement is reflected in rhythmic neural activity in the alpha (~ 8–12 Hz) and beta (~ 14–30 Hz) frequency range in sensory and higher-order brain areas (e.g., Ahveninen, Huang, Belliveau, Chang, & Hämäläinen, 2013; Deiber, Ibañez, Missonnier, Rodriguez, & Giannakopoulos, 2013; Frey, Ruhnau, & Weisz, 2015; Hanslmayr, Staudigl, & Fellner, 2012; Jensen & Mazaheri, 2010; Klimesch, Sauseng, & Hanslmayr, 2007; Sander, Werkle-Bergner, & Lindenberger, 2012; Waldhauser, Johansson, & Hanslmayr, 2012). Generally, it is assumed that alpha rhythmic activity routes task-relevant information by inhibiting information processing in task-irrelevant brain areas (Jensen & Mazaheri, 2010; Klimesch et al., 2007). Effective inhibition of task-irrelevant regions is thereby indicated by increased alpha power (i.e., alpha synchronization). Active processing, in turn, is accompanied by a decrease in alpha rhythmic activity (i.e., alpha de-synchronization; e.g., Hanslmayr, Staudigl, & Fellner, 2012; Waldhauser, Johansson, & Hanslmayr, 2012; Werkle-Bergner et al., 2014).

Typically, during visuospatial selective attention tasks, alpha rhythmic activity in parieto-occipital regions is lateralized, with enhanced alpha-power over the hemisphere ipsilateral to the attentional focus and contralateral to the irrelevant distractor, whereas alpha power is reduced contralateral to the target item (Kerlin, Shahin, & Miller, 2010; cf. Figure 1, bottom row). This pattern has also been demonstrated during the pre-stimulus phase after the presentation of a spatial attention cue (e.g., Bauer, Stenner, Friston, & Dolan, 2014; for an overview, see Jensen & Mazaheri, 2010) indicating the deployment of spatially selective attention already during stimulus anticipation – a largely internally driven process.

Comparable effects were also found in auditory tasks in the alpha band (Ahveninen et al., 2013; Banerjee, Snyder, Molholm, & Foxe, 2011; Frey et al., 2014; Wöstmann, Herrmann, Maess, & Obleser, 2016) and for somatosensory and visuomotor selective attention tasks (e.g., Bauer, Kennett, & Driver, 2012; see Engel & Fries for overview, 2010; Gola, Magnuski, Szumska, & Wróbel, 2013) mainly in the beta band. In sum, across modalities spatially specific modulations of ongoing rhythmic neural activity in the alpha–beta range (~ 8–30 Hz) indicate the spatially specific deployment of attention, reflecting either the suppression of distracting sensory input, facilitated processing of target information or both (for an overview, see Frey, Ruhnau, & Weisz, 2015).

Studies employing attentional and working memory (WM) tasks suggest altered alpha–beta rhythmic activity across the adult lifespan (e.g., Deiber et al., 2013; Gola, Kamiński, Brzezicka, & Wróbel, 2012; Gola et al., 2013; Hong, Sun, Bengson, Mangun, & Tong, 2015; Leenders, Lozano-Soldevilla, Roberts, Jensen, & De Weerd, 2018; Rogers, Payne, Maharjan, Wingfield, & Sekuler, 2018; Sander et al., 2012). Thereby, the majority of studies report decreased or even absent posterior-parietal alpha–beta-lateralization in older adults. For example, alpha lateralization has been shown to be absent in older compared to younger adults in a visuospatial selective attention task (Hong et al., 2015) and during retention in a visual WM task (Leenders et al., 2018; Sander et al., 2012). However, during low load retention (Sander et al., 2012) and attentional cueing (Leenders et al., 2018; Mok, Myers, Wallis, & Nobre, 2016) within WM alpha lateralization was preserved. Thus, selective attention mechanisms reflected in posterior alpha lateralization in visual tasks do not seem to be completely compromised by normal aging. Rather the magnitude of the age differences depends on the specific task demands. Furthermore, in the absence of age-related differences in task performance older compared to younger adults showed an increase in central, sensorimotor beta activity, while at the same time attention-related alpha rhythmic activity in posterior areas was decreased (e.g., Deiber et al., 2013) reflecting a functional reorganization of rhythmic neural acitvity with increasing age.

Studies investigating age-related changes in alpha lateralization in the auditory domain are scarce. To the best of our knowledge, there is, so far, only one very recent study investigating age differences in alpha lateralization during a trial-by-trial cued auditory spatial selective attention task (Tune, Wöstmann, & Obleser, 2018). This study indicated a youth-like alpha lateralization (8–12 Hz) in a sample of high performing middle-aged to older adults. Given that the performance of the studied, rather young older-adult sample (~ 40–70 years) was comparable to younger adults tested in a previous study (Wöstmann et al., 2016), it is still an open question whether the alpha lateralization effect would be present in a sample of older adults with declined spatial selective attention abilities.

Thus, the aim of the present study was to examine age-related differences in rhythmic neural activity in the alpha–beta (~ 8–30 Hz) frequency range during auditory spatial selective attention. Specifically, we were interested in rhythmic neural signatures of self-initiated, preparatory (i.e., pre-stimulus alpha-lateralization) and externally driven, reactive control over competing auditory stimuli (i.e., post-stimulus alpha–beta lateralization).

For this purpose, we re-analyzed EEG data of a previously published event-related potential study (Passow et al., 2014) in which samples of younger (22–35 years) and older adults (63–76 years) were tested in an attention and intensity-modulated dichotic listening task (cf. Westerhausen et al., 2009; see Methods). In this task, two streams of highly similar auditory input were simultaneously presented to participants′ both ears (i.e., dichotically) while attention had to be focused on the input to only one ear (i.e. target) and the other, distracting information had to be ignored. In the light of previous work (e.g., Ahveninen et al., 2013; Banerjee et al., 2011; Frey et al., 2014; Weisz, Hartmann, Müller, Lorenz, & Obleser, 2011; Wöstmann et al., 2016) we expected younger adults to exhibit lateralized alpha-band activity, i.e. an alpha synchronization ipsilateral and an alpha desynchronization contralateral to the target ear, before and after stimulus presentation (see Figure 1, bottom row). We hypothesized this effect to be more pronounced for correct compared to incorrect trials, reflecting self-initiated attentional enhancement of auditory target signals (e.g., Tune et al., 2018; Wöstmann et al., 2016; see Figure 1, middle row). Given the reported age-related decreases in alpha lateralization in visual selective attention tasks (Hong et al., 2015; Leenders et al., 2018; Sander et al., 2012), we expected the alpha lateralization effect to be much weaker or even absent in older adults (but see Tune et al., 2018 for different observations; see Figure 1, top row). Importantly, in line with an age-graded shift from internal to externally driven attentional control (Lindenberger & Mayr, 2014), we expected diminished alpha-lateralization in older adults mainly before stimulus onset (i.e., in an internally driven, anticipatory phase). Finally, to demonstrate the behavioral relevance of moment-to-moment adaptations in lateralized rhythmic neural activity we conducted single trial (mixed-effects logistic regression) analyses.

**Figure 1.**
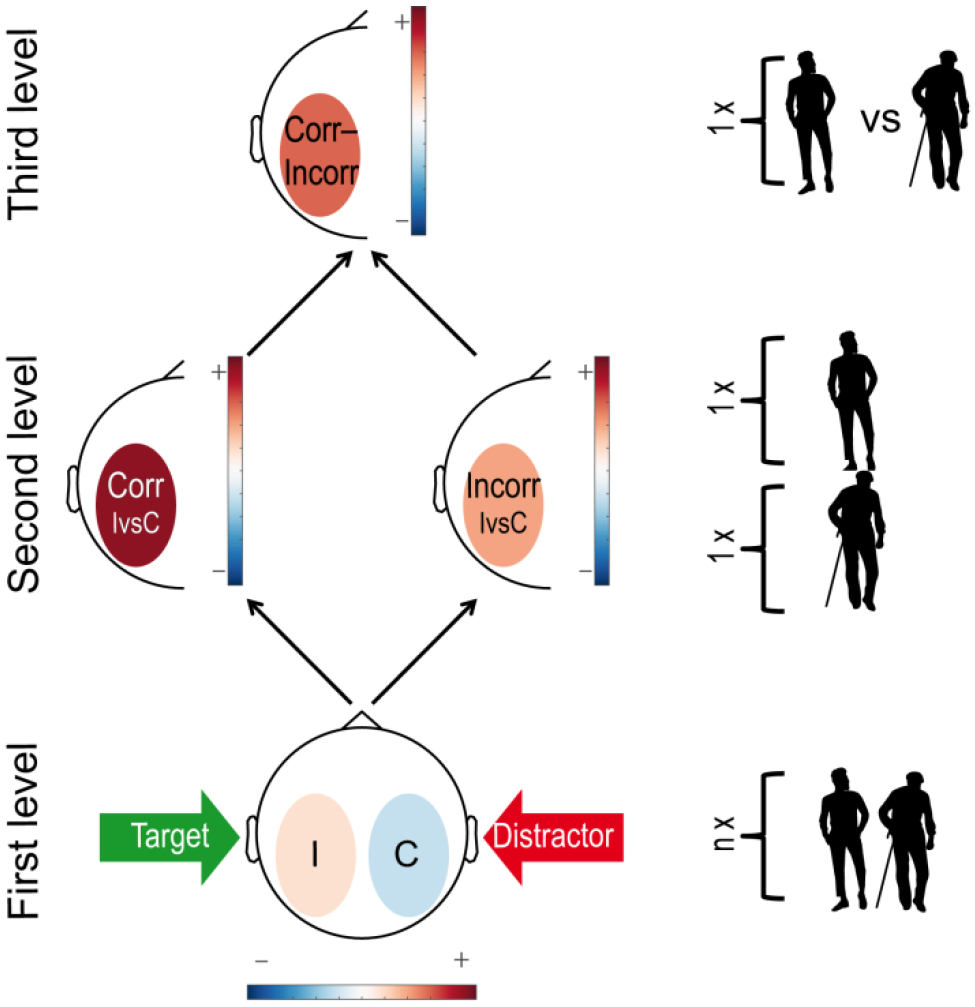
Schematic overview over levels of EEG data analysis in relation to the hypotheses we tested. Within subjects (first level), ipsi (I)- and contralateral (C) power, reflecting the processing of distractor and target processing, respectively, is contrasted to identify task-related lateralized patterns of rhythmic neural activity. On a second level (within age groups), lateralized activity of correct and incorrect trials is compared to identify age group specific rhythmic neural correlates associated with successful selective attention performance. On a third level (across age groups), rhythmic neural correlates of correct performance (as identified on a second level) were investigated for age differences.

## Results

### Behavioral results

#### Auditory selective attention is reduced in older adults

In the dichotic listening task, participants successfully adjusted their attentional focus to relevant stimuli (i.e., according to cue condition) as reflected by a main effect of Attentional Focus: *F*(1.488, 72.924) = 32.748, *p* < 0.001, *η*² = 0.401 resulting from a two-factorial repeated measures ANOVA (Age group × Attentional Focus). The Age group main effect did not reach significance (*F*(1, 49) = 1.359, *p* = 0.249, *η*² = 0.027); however, younger and older adults differed reliably in their ability to modulate their focus as indicated by a significant Age group × Attentional focus interaction (*F*(1.488, 72.924) = 10.659, *p* < 0.001, *η*² = 0.179; see Figure 2).

While demonstrating reliably different levels of selective attention (i.e., a significant Age group × Attentional Focus interaction), post-hoc tests indicated successful selective attention within both age groups (younger adults: *F*(1.183, 28.382) = 30.829, *p* < 0.001, *η*² = 0.562; older adults: *F*(1.833, 45.822) = 6.043, *p* = 0.006, *η*² = 0.195).

**Figure 2.**
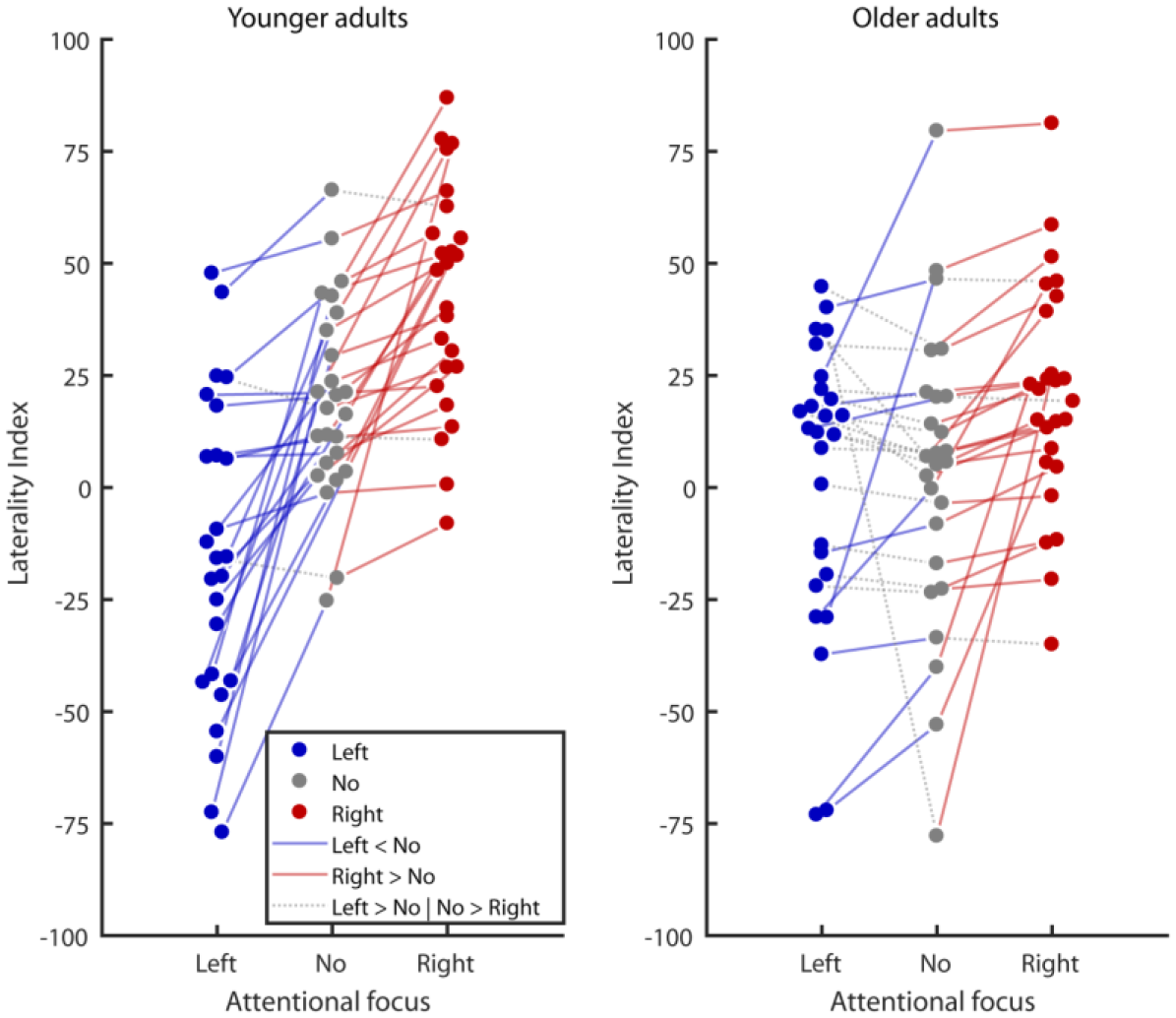
Selective attention performance of younger (YA) and older adults (OA) in a dichotic listening task. Negative and positive laterality index values indicate a tendency for left and right ear responses, respectively. Circles connected by solid colored lines show participants who demonstrate a behavioral selective attention effect (i.e., more responses of the cued ear relative to the No focus condition), with the slope of the lines reflecting the degrees of selective attention. Circles connected by grey dotted lines indicate a reversed effect. While the amount of selective attention is markedly decreased in older adults, both younger and older participants demonstrate reliable selective attention on a group level.

### Electrophysiological results

#### Different pre- and post-stimulus lateralization patterns of rhythmic neural activity relate to successful selective attention within younger and older adults

In younger adults, successful selective attention performance was associated with sustained lateralized rhythmic neural activity in an extended alpha frequency range (6–16 Hz) spanning pre- and post-stimulus time periods (−0.828–0.890 s with respect to stimulus onset; see Figure 3). The observed cluster (*p*_*corr*_ = 0.021) consisted mainly of parieto-occipital electrodes (see Figure 3).

**Figure 3.**
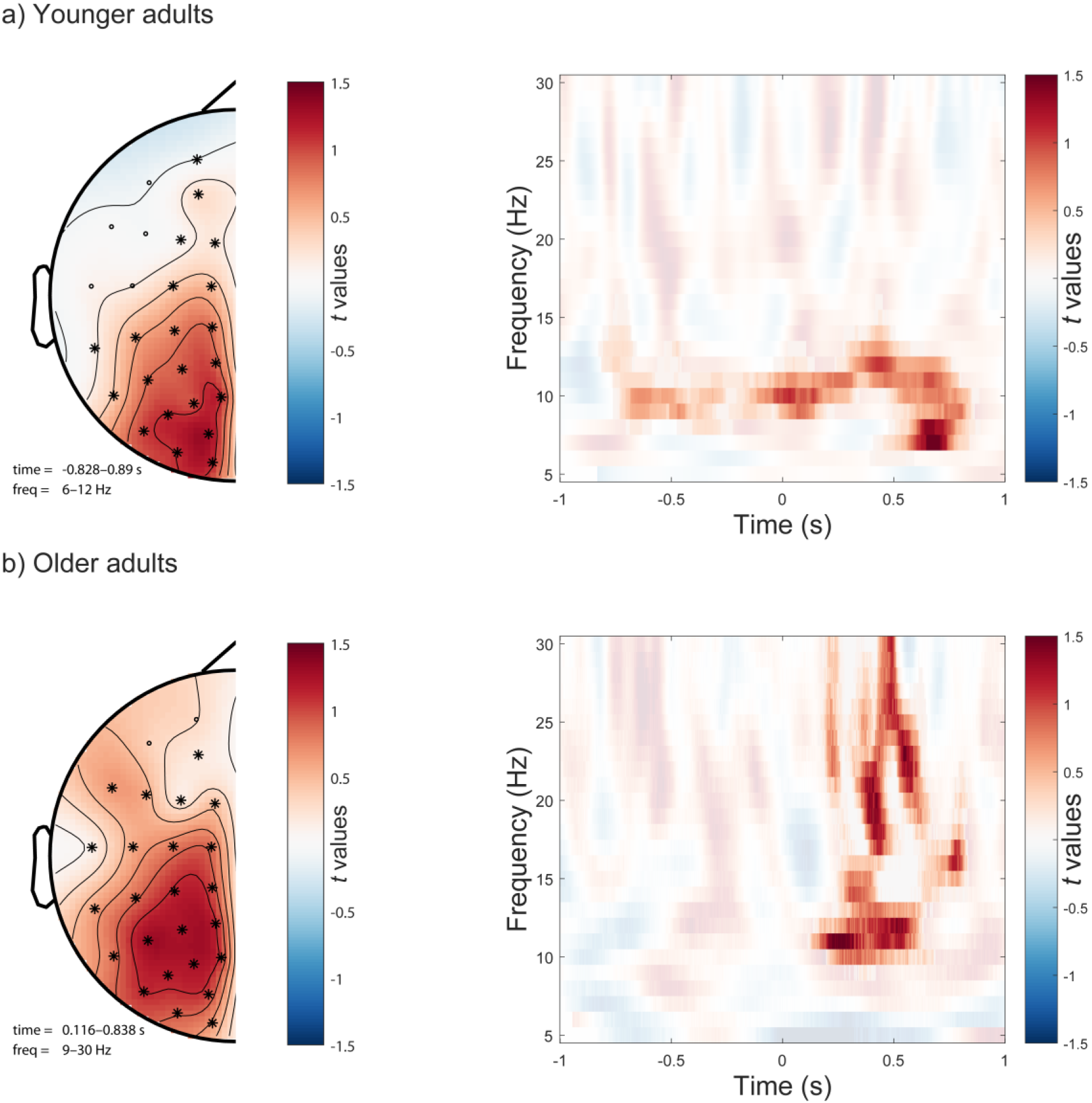
Electrodes, and time-frequency samples associated with correct selective attention performance in younger (a) and older (b) adults. (left panel, a): Mainly parieto-occipital electrodes constitute the significant cluster in younger adults. (right panel, a): Sustained (pre- and post-stimulus) alpha-activity is associated with correct performance in younger adults. Semitransparent time-frequency samples are not part of the significant cluster. (Left panel, b): Mainly centro-parietal electrodes constitute the significant cluster in older adults. (Right panel, b): Only post-stimulus alpha–beta-activity is associated with correct performance in older adults. Semitransparent time-frequency samples are not part of the significant cluster.

In older adults, correct performance was related to activity in the alpha–beta frequency range (9–30 Hz) that was, however, restricted only to the post-stimulus time period (0.116–0.838 s with respect to stimulus onset; see Figure 3). The older adults′ cluster (*p*_*corr*_ = 0.001) also included mainly parieto-occipital electrodes but was shifted more laterally compared to the younger adult′s cluster (see Figure 3).

Significant cluster results were followed up by means of two-factorial (Ipsi/Contra × Correct/Incorrect) ANOVAs within each age group. In younger adults, we did not observe reliable differences in lateralized alpha activity (6–16 Hz) related to performance (main effect Correct/Incorrect: *F*(1, 24) = 0.024, *p* = 0.877, *η*² = 0.001) or hemisphere (Ipsi/Contra: *F*(1, 24) = 3.996, *p* = 0.057, *η*² = 0.143). Crucially, however, we detected a reliable performance × hemisphere interaction (Correct/Incorrect × Ipsi/Contra: *F*(1, 24) = 6.826, *p* = 0.015, *η*² = 0.221), indicating higher ipsi- and lower contralateral power in correct compared to incorrect trials, respectively (see Figure 4 a).

In older adults, post-hoc analyses also did not reveal reliable differences in lateralized alpha–beta activity (9–30 Hz) related to performance (main effect Correct/Incorrect: *F*(1, 25) = 1.037, *p* = 0.318, *η*² = 0.040) or hemisphere (Ipsi/Contra: *F*(1, 25) = 0.597, *p* = 0.447, *η*² = 0.023). While numerically in the same direction as in younger adults, in older adults the performance × hemisphere interaction, however, failed to reach significance (Correct/Incorrect × Ipsi/Contra: *F*(1, 25) = 2.496, *p* = 0.127, *η*² = 0.091) (see Figure 4 b).

**Figure 4.**
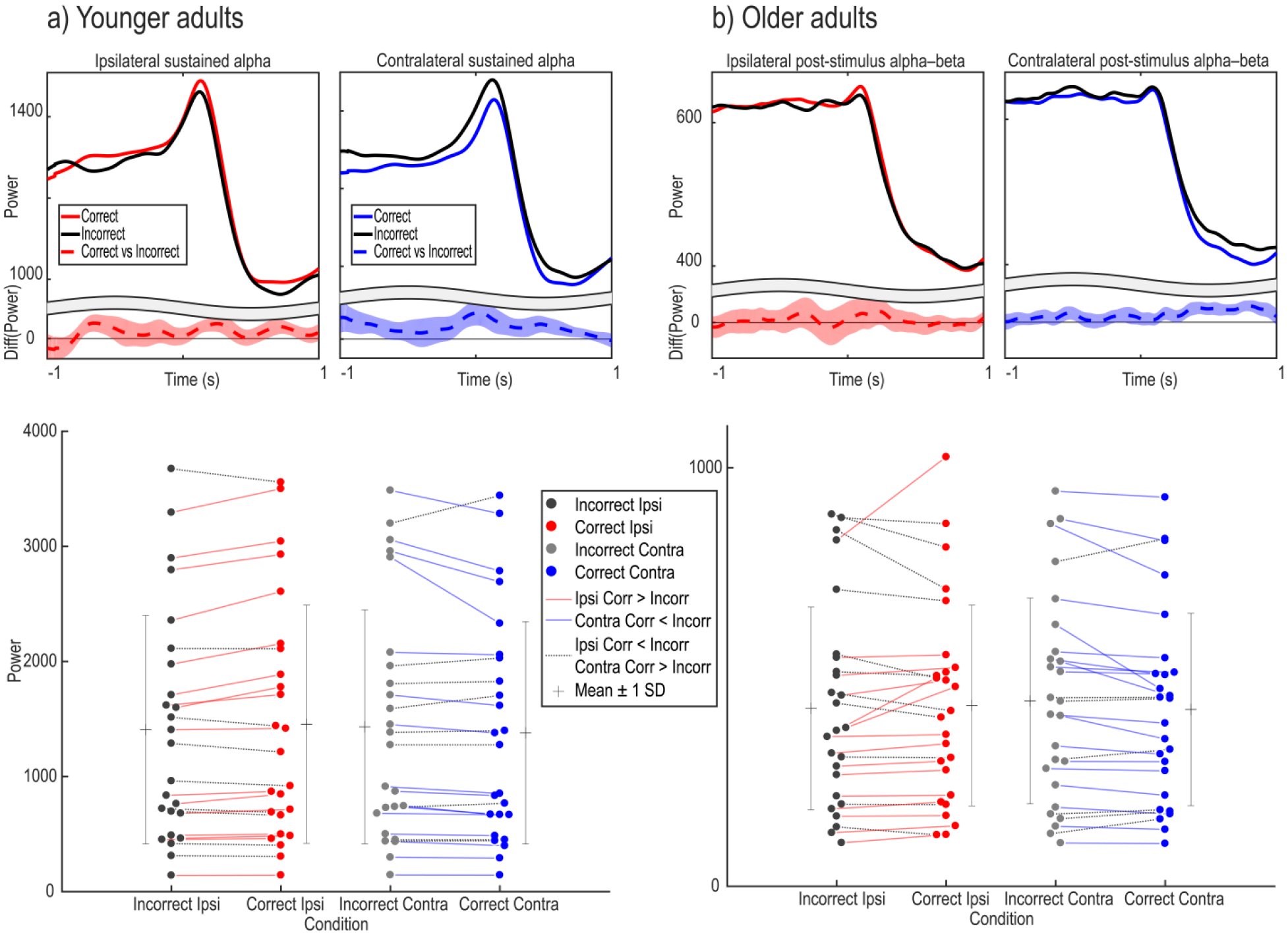
(a) Younger adults′ ipsi- and contralateral sustained alpha (6–16 Hz) activity split for correct and incorrect trials. (b) Older adults′ ipsi- and contralateral post-stimulus alpha–beta (9–30 Hz) activity split for correct and incorrect trials. (Upper row): Time course (± 1 s with respect to stimulus onset) of ipsi- and contralateral de/synchronization for correct (solid colored lines) and incorrect (solid black lines) trials. Broken lines indicate the ipsi vs. contra difference for each time point. Shaded areas show ± 1 standard error of the mean. (Lower row): A two-factorial (Ipsi/Contra × Correct/Incorrect) ANOVA reveals a reliable interaction effect (i.e., higher ipsilateral and lower contralateral power for correct relative to incorrect trials) in YA but not OA. Circles connected by colored lines show participants that demonstrate the hypothesized interaction effect while dotted black lines indicate a reversed effect. The plus signs show the means (per condition and age group) ± 1 standard deviation.

#### Diminished pre-stimulus alpha-lateralization in older adults

After identifying rhythmic neural correlates of correct selective attention performance within each age group, we investigated whether these differed reliably across groups. We observed robust age-group differences in the extended alpha frequency range (6–12 Hz) spanning pre- and post-stimulus time periods (*U*(49) = 758; *Z* = 2.026; *p* = 0.043). Crucially, post-hoc tests indicated that this age difference was driven by lower pre-stimulus alpha-lateralization in older adults (pre-stimulus: *U*(49) = 814; *Z* = 3.081; *p* = 0.002; post-stimulus: *U*(49) = 726; *Z* = 1.423; *p* = 0.155; see Figure 5). Older adults in turn exhibited stronger post-stimulus alpha–beta activity (9–30 Hz) relative to younger adults (*U*(49) = 460; *Z* = −3.571; *p* < 0.001; see Figure 5). In sum, older adults demonstrated reduced preparatory alpha-lateralization (i.e., before stimulus onset) and increased alpha–beta lateralization in response to stimulus presentation, in line with an age-graded shift from internal to externally driven control (Lindenberger & Mayr, 2014).

**Figure 5.**
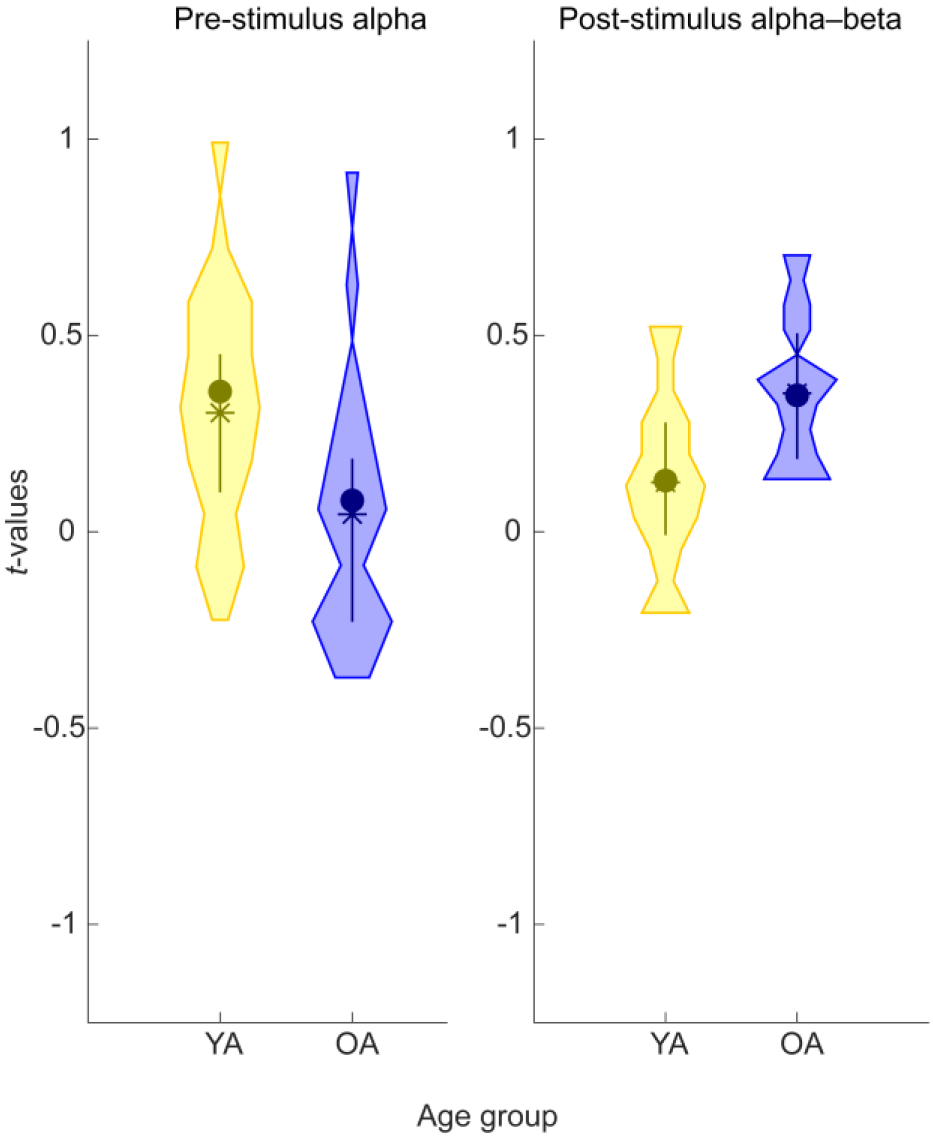
Age differences in rhythmic neural correlates of successful selective attention between younger (YA; yellow) and older (OA; blue) adults during pre- and post-stimulus time windows. Higher *t*-values (y-axis) indicate stronger lateralized rhythmic neural activity during successful relative to unsuccessful selective attention. Left panel: Older adults show lower pre-stimulus alpha (6–12 Hz) activity relative to younger adults (*p* = 0.002). Right panel: Older adults demonstrate significantly stronger post-stimulus alpha–beta (9–30 Hz) activity compared to younger adults (*p* < 0.001).

#### Age group specific prediction of single trial accuracy by indicators of lateralized rhythmic neural activity

The behavioral relevance of moment-to-moment fluctuations in lateralized rhythmic neural activity as well as potential age differences therein were evaluated by means of mixed-effects logistic regression analyses. In particular, the accuracy in each trial was predicted by its sustained alpha (see YA cluster, Figure 3) and post-stimulus alpha–beta (see OA cluster, Figure 3) activity as well as trial condition and age group. To investigate age-group specific effects of lateralized rhythmic neural activity, we additionally included (1) Age group × SustainedAlpha and (2) Age group × Post-stimulusAlpha–beta interaction terms. Individual performance differences were modeled as random-effects of intercept grouped by individual participant (ID). The model formula in Wilkinson notation is expressed below.

*Equation 1:*

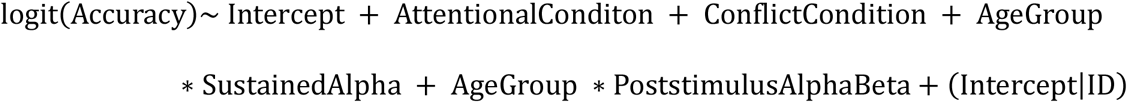

The full model (see Eq. 1 and Figure 6) fit the data well (Log Likelihood: −16034; AIC: 32088; BIC: 32170; adjusted *r*^2^ = 0.183) and significantly outperformed a constant only model (LR(9) = 3831.2; *p* < 0.001). Further, the model showed reliably better fit to the data compared to a model without EEG predictors (i.e., SustainedAlpha and PoststimulusAlphaBeta; LR(4) = 62.704; *p* < 0.001). Finally, we compared the full model to a fixed-effects only model including otherwise the same factors, indicating reliable interindividual differences (LR(1) = 1102.8; *p* < 0.001).

**Figure 6.**
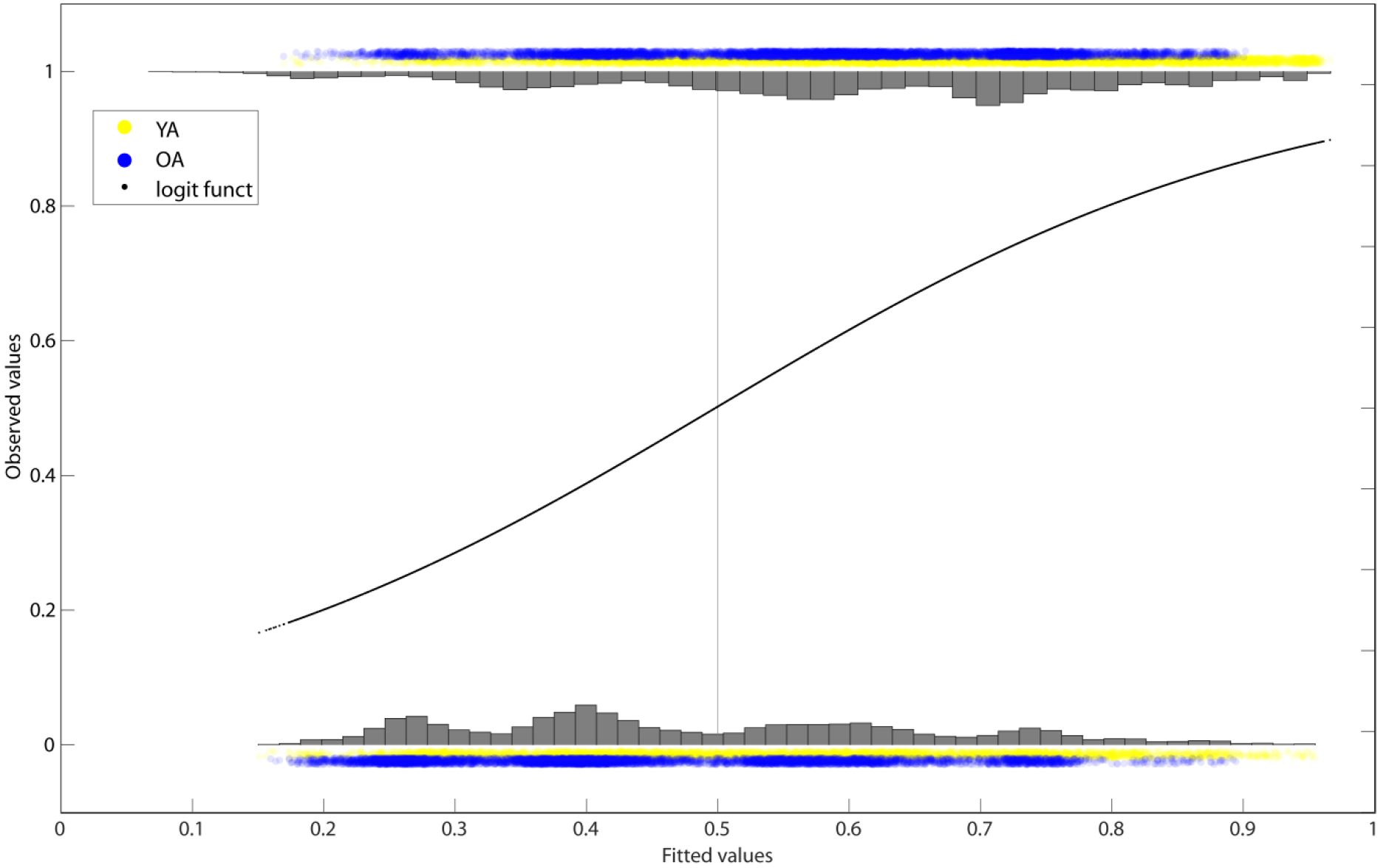
Single trial logistic regression model-predicted (fitted, x-axis) and actually observed accuracy values (y-axis) for younger (YA; yellow) and older (OA; blue) adults. Fitted values below/above 0.5 (grey line) indicate a prediction of in/correct performance, respectively. As evident from the skewed histograms (i.e. not centered around 0.5), the model demonstrates above-chance prediction accuracy of single trial performance. This is, higher model-predicted accuracy values coincide with actually observed correct performance and vice versa.

Single trial lateralized rhythmic neural activity was robustly linked to performance. Across younger and older subjects, higher sustained alpha-lateralization was reliably associated with better task performance (*t*(26231) = 6.268; *p* < 0.001; standardized estimate: 0.160; see Table 1 and Figure 7), whereas post-stimulus alpha–beta lateralization was not (*t*(26231) = −1.274; *p* = 0.203; see Table 1 and Figure 7). Crucially, we observed reliable Age group × SustainedAlpha and Age group × Post-stimulusAlpha–beta interactions (see Table 1 and Figure 7). In older adults, higher sustained alpha lateralization was related to worse performance (*t*(26231) = −6.443; *p* < 0.001; standardized estimate: −0.218). In contrast, higher post-stimulus alpha–beta lateralization was associated with better performance in old age (*t*(26231) = 4.016; *p* < 0.001; standardized estimate: 0.130). That is, sustained alpha-lateralization and post-stimulus alpha–beta lateralization showed an age-group specific association to selective attention performance on a single trial level. While in younger adults higher sustained alpha-lateralization increased performance, in older adults it had detrimental effects on accuracy. Similarly, post-stimulus alpha–beta lateralization improved accuracy only in older adults and was not linked to behavior in younger adults. Of note, a highly comparable pattern of results emerged when restricting single trial estimates of alpha-activity to the pre-stimulus time period (PreStimulusAlpha *t*(26231) = 3.813; *p* < 0.001; standardized estimate: 0.087; PreStimulusAlpha * OldAge *t*(26231) = −5.436; *p* < 0.001; standardized estimate: −0.168).

**Table 1.** Fixed effect coefficients for single trial logistic regression analyses

| Fixed effects | Estimate | Standard error | *t*-value | *df* | *p* |
| --- | --- | --- | --- | --- | --- |
| Intercept | 1.085 | 0.105 | 10.345 | 26231 | < 0.001 |
| OldAge | −0.531 | 0.143 | −3.725 | 26231 | < 0.001 |
| SustainedAlpha | 0.160 | 0.026 | 6.268 | 26231 | < 0.001 |
| PoststimulusAlphaBeta | −0.031 | 0.024 | −1.274 | 26231 | 0.203 |
| SustainedAlpha * OldAge | −0.218 | 0.034 | −6.443 | 26231 | < 0.001 |
| PoststimulusAlphaBeta * OldAge | 0.130 | 0.032 | 4.016 | 26231 | < 0.001 |
| AttentionalCondition (FR) | 0.536 | 0.027 | 19.791 | 26231 | < 0.001 |
| ConflictCondition (no) | −0.807 | 0.034 | −24.053 | 26231 | < 0.001 |
| ConflictCondition (high) | −1.488 | 0.034 | −43.660 | 26231 | < 0.001 |

*Note:* Model formula is expressed in Equation 1.

**Table 2.** Perceptual speed and verbal knowledge for younger and older adults

| Measure | Younger adults | Older adults | *t*-value | *df* | *p* | *d* |
| --- | --- | --- | --- | --- | --- | --- |
| Digit Symbol Substitution test | $71.5 \pm 13.9$ | $48.3 \pm 11.3$ | 6.54 | 49 | $< 0.001$ | 1.83 |
| Spot-A-Word | $18.2 \pm 5.3$ | $23.7 \pm 4.3$ | -4.02 | 49 | $< 0.001$ | 1.12 |

Note: For younger and older adults, group mean performance $\pm 1$ standard error of the mean is reported.

**Figure 7.**
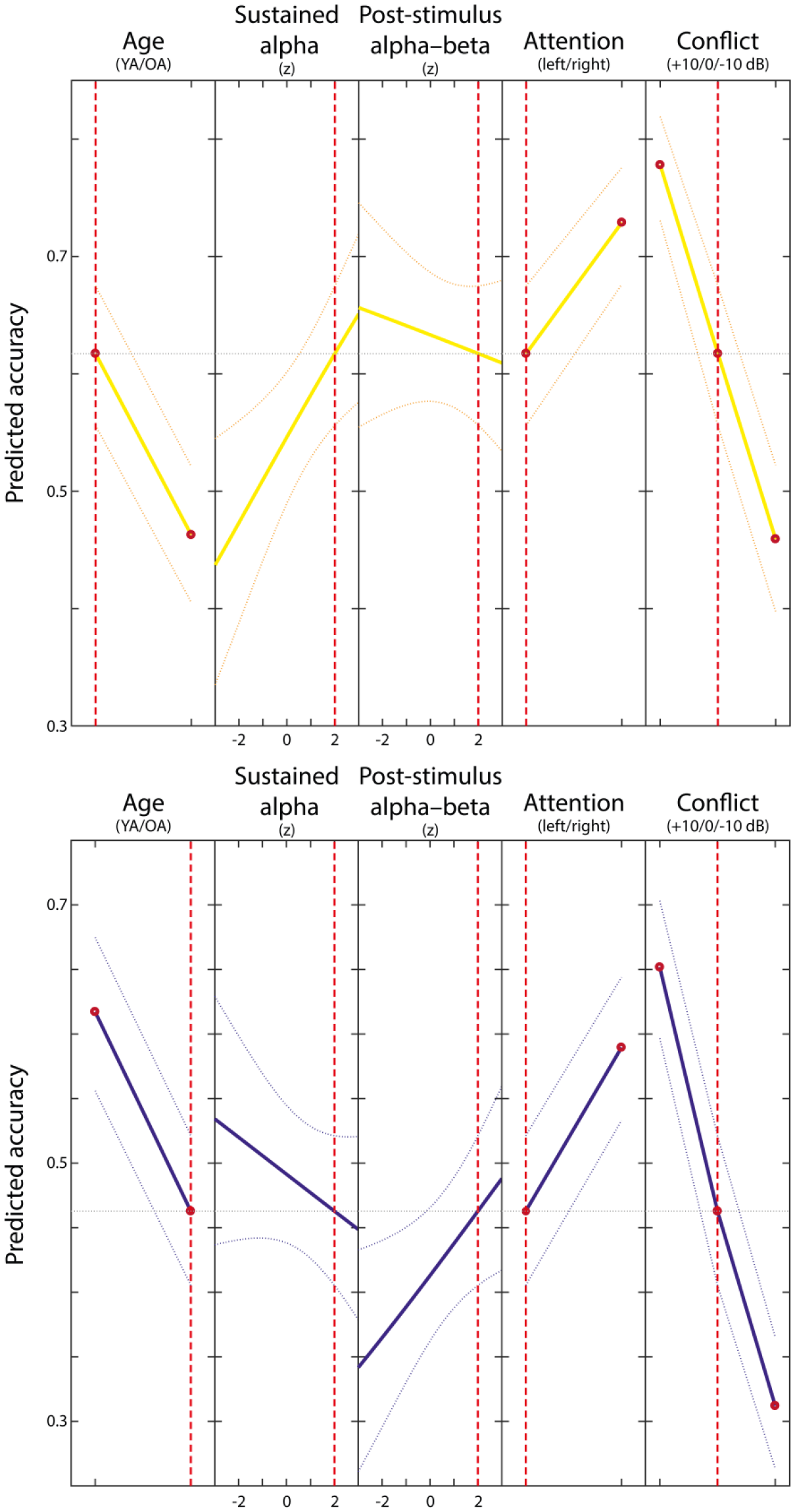
Plot of slices through fitted logistic regression surface for younger (upper plot; yellow) and older (lower plot; blue) adults. Dotted yellow/blue lines show ± 1 standard error of the mean. Red broken vertical lines indicate the depicted conditions. While in younger adults higher sustained alpha-lateralization is associated with accuracy, in older adults higher post-stimulus alpha–beta lateralization is linked to single trial performance. Old age, left-cued (FL) trials and high perceptual conflict (i.e., [−10] in case of FR; [+10] in case of FL) are predictive of lower accuracy. For visualization, a model without random-effects is depicted.

Across age groups, performance was higher in trials in which attention was cued to the right ear which reflects the well-established right ear advantage of verbal auditory processing (see Hugdahl, 2003; Kimura, 1967 for reviews) and when no or only low perceptual conflict was present (see Table 1 and Figure 7).

## Discussion

Older adults experience difficulties in daily situations requiring flexible information selection in the presence of multiple competing sensory inputs, like for instance multi-talker situations (e.g., Hugdahl et al., 2009; Passow et al., 2012, 2014; Westerhausen, Bless, & Kompus, 2015). Modulations of rhythmic neural activity in the alpha–beta (8–30 Hz) frequency range in sensory and higher-order brain areas have been established as a cross-modal neural correlate of selective attention (e.g., Frey et al., 2015; Hanslmayr et al., 2012; Jensen & Mazaheri, 2010; Klimesch et al., 2007). However, research linking compromised auditory selective attention to changes in rhythmic neural activity in aging is sparse (for a notable exception, see Tune, Wöstmann, & Obleser, 2018).

Thus, this study investigated adult age differences in rhythmic neural activity in the alpha–beta frequency range (~ 8–30 Hz) during auditory spatial selective attention. Compared to younger adults, older adults′ performance was severely compromised in this task. On the neurophysiological level, we observed lateralized rhythmic neural activity in an extended alpha frequency range (6–16 Hz) during the anticipatory, pre-stimulus phase in younger but not older adults. After stimulus presentation, alpha lateralization was sustained in younger adults and a spatially distinct, rhythmic neural activity pattern in the alpha–beta frequency (9–30 Hz) range emerged in older adults. Importantly, these age group specific rhythmic neural activity patterns were significantly more pronounced during successful (relative to unsuccessful) selective attention, underlining their behavioral relevance. Single trial analyses investigating the functional significance of moment-to-moment fluctuations in rhythmic neural activity further supported an age group specific association between pre-and post-stimulus activity and selective attention. In particular, we observed that higher preparatory, pre-stimulus alpha rhythmic activity was associated with better performance in younger but lower accuracy in older adults. In contrast, higher stimulus-driven alpha–beta lateralization predicted more accurate behavior in older adults but was unrelated to performance in younger adults. In the following we will discuss these results in more detail.

On the behavioral level, both, younger and older adults were able to modulate their attention flexibly according to changing task demands as assessed using a dichotic listening paradigm (cf. Westerhausen et al., 2009). However, in line with earlier findings, older adults showed a severe decline in attentional control of auditory perception compared to younger adults (e.g., Passow et al., 2012, 2014; Westerhausen, Bless, Passow, Kompus, & Hugdahl, 2015).

On the neural level, younger adults demonstrated a sustained pattern of lateralized rhythmic neural activity in an extended alpha (6–16 Hz) frequency range at mainly parietooccipital electrodes as previously reported (e.g., Ahveninen et al., 2013; Banerjee et al., 2011; Frey et al., 2014; Wöstmann et al., 2016). Follow-up analyses further revealed that this effect was driven by alpha-synchronization in currently task irrelevant (i.e., ipsilateral) and alpha-desynchronization in task relevant (i.e., contralateral) brain areas. This indicates that in younger adults successful selective attention was promoted by the active inhibition and facilitation of distractor and target processing, respectively (Frey et al., 2015; Jensen & Mazaheri, 2010; Klimesch et al., 2007; Weisz et al., 2011). Importantly, in the employed dichotic listening task participants′ attentional focus was cued by a single attentional instruction at the beginning of each block (i.e., once every ~ 80 trials, without trial-by-trial changes of attentional focus), allowing participants to prepare for upcoming stimuli before their actual presentation. Younger adults utilized this information and modulated the balance of cortical excitation and inhibition (i.e., alpha de/synchronization) in sensory areas according to task demands, reaching statistical significance as early as 800 ms before actual stimulus presentation (Jensen & Mazaheri, 2010; Klimesch et al., 2007). Of note, in the absence of an external attentional cue this indicates internally driven, self-initiated attentional control in younger adults (Craik & Bialystok, 2006; Lindenberger & Mayr, 2014) In contrast, successful selective attention in older adults was associated with merely stimulus-driven, lateralized rhythmic neural activity in the alpha–beta (9–30 Hz) frequency range. This is, effective lateralization of rhythmic neural activity reflecting the flexible routing of information selection depended on external input (i.e., stimulus presentation) in older adults. Our findings are thus in line with an age-graded shift from self-initiated towards externally driven control of attentional processing (Craik & Bialystok, 2006; Lindenberger & Mayr, 2014). Previous visuospatial selective attention studies reported age-related decreases in alpha modulation (e.g., Deiber, Ibañez, Missonnier, Rodriguez, & Giannakopoulos, 2013; Hong et al., 2015; Leenders et al., 2018; Sander et al., 2012). In particular, employing an adapted Attentional-Network Task (Fan, McCandliss, Sommer, Raz, & Posner, 2002) Deiber and colleagues (2013) investigated age differences in rhythmic neural activity related to anticipatory and stimulus-driven attentional processes. They observed less anticipatory alpha modulation in older adults, especially when the demand for self-initiated processing was high (i.e., when no external attention cue predicted the target occurrence; Deiber et al., 2013). After presentation of the target stimulus, however, older adults exhibited a reliable modulation of alpha–beta activity that was shifted to higher frequencies and more central electrodes compared to younger adults′ response. Deiber and colleagues (2013) interpreted their observation as support for a partial reorganization of attention-related rhythmic neural responses with age, consisting of a decreased activation of posterior, attentional resources and an additional recruitment of central, sensorimotor circuits. The diminished anticipatory alpha lateralization we report here closely lines up with these findings and moreover suggests a cross-modal age-related impairment in self-initiated selective attention (cf. Deiber et al., 2013; Hong et al., 2015).

Critically, our study complements previous work by demonstrating age differences in the functional significance of moment-to-moment fluctuations in rhythmic neural activity even on a single trial level. Successful selective attention was predicted by higher anticipatory alpha lateralization in younger adults. In contrast, pre-stimulus alpha rhythmic neural activity was associated with significantly worse performance in older adults. Stimulus-driven alpha–beta lateralization in turn improved accuracy only in older adults and was not linked to behavior in younger adults. These findings further support an age-related reorganization of attentional processing, including a shift from internal, preparatory to external, stimulus-driven control (Craik & Bialystok, 2006; Lindenberger & Mayr, 2014) evident in central alpha–beta rhythmic activity (Deiber et al., 2013; Gola et al., 2012, 2013). Higher preparatory alpha lateralization reduced performance in older adults suggesting older adults′ shift as a strategic adaptation to diminishing attentional resources with age (Velanova et al., 2006). In particular, previous research demonstrated a reduced ability to maintain heightened level of alpha modulation over longer periods of time in old age (Wöstmann, Herrmann, Wilsch, & Obleser, 2015), especially in difficult task conditions (Sander et al., 2012). Thus, older adults′ shift to stimulus-driven attentional control may pose a strategic adaption to prevent a premature breakdown of alpha lateralization before actual stimulus presentation. In line with earlier reports, single trial performance in younger and older adults was higher in trials in which attention was cued to the right ear (Hugdahl et al., 2009) and when no or only low perceptual-attentional conflict was present (Passow et al., 2012, 2014; Westerhausen et al., 2009).

In contrast to our finding of age differences in selective attention and alpha lateralization, a very recent study reported a well-preserved alpha modulation in a sample of middle-aged to older adults during a trial-by-trial cued auditory spatial selective attention task (Tune et al., 2018). A number of reasons may explain the observed discrepancy. First, as highlighted above, older adults′ ability to flexibly adapt alpha rhythmic activity critically depends on task difficulty (Sander et al., 2012), breaking down when faced with excessive demands. The overall younger sample (aged 39–69 years; n = 23) tested by Tune and colleagues (2018), however, performed on a high, youth-like level (cf. Wöstmann et al., 2016) indicating low–moderate task difficulty. In contrast, while both younger and older adults were able to exert selective attention, we observed severely compromised performance in older adults implying high difficulty. Second, in line with an age-graded shift from internal to external control (Lindenberger & Mayr, 2014) age differences in the modulation of alpha rhythmic neural activity appear particularly exacerbated in absence of external stimuli (cf. Deiber et al., 2013). While Tune and colleagues (2018) supported older adults with a spatial attention cue on each trial, our participants had to exert purely self-initiated control, probably contributing to the heightened task difficulty (Craik & Bialystok, 2006). Finally, Tune and colleagues (2018) restricted their analyses a-priori to a cortical region and frequency of interest (i.e., alpha at posterior electrodes), potentially obscuring an age-specific rhythmic neural response shifted in space and frequency. In sum, together with the present results this indicates that in aging the integrity of selective attention and its neural implementation is moderated by additional factors, like task difficulty.

Recently, also Rogers and colleagues reported age differences in alpha rhythmic activity during auditory attention (Rogers et al., 2018). A direct comparison of results, however, is impeded by two important aspects. First, despite an indication for age-related hearing loss in their older adult sample, no frequency-specific adjustment of stimulus volumes (cf. Passow et al., 2012, 2014; Tune et al., 2018; Wöstmann et al., 2015) was applied, complicating the distinction between perceptual and attentional age differences (Li, Lindenberger, & Sikstrom, 2001). Second and more crucially, unlike previous work, age comparisons were not calculated on neural measures of spatially selective attention (e.g., alpha-modulation index; AMI; or lateralized rhythmic neural activity; cf. Sander et al., 2012; Tune et al., 2018; Wöstmann et al., 2016). Instead age differences are only reported for neural correlates of auditory stimulus processing irrespective of focused selective attention (i.e., averaged across focus left and focus right conditions), precluding direct comparisons.

To conclude, selective attention is required to flexibly route information in the presence of multiple competing sensory inputs, like entailed in daily social interactions. We here demonstrate a link between severely compromised auditory selective attention and a partial reorganization of attention-related rhythmic neural responses in aging. In particular, in old age we observed a shift from a self-initiated, preparatory modulation of alpha rhythmic activity to an externally driven, response in the alpha–beta range. Critically, moment-to-moment fluctuations in the age-specific patterns of self-initiated and externally driven lateralized rhythmic activity were tied to selective attention. We conclude that adult age difference in spatial selective attention likely derive from a functional reorganization of rhythmic neural activity within the aging brain.

## Methods

### Participants

In the present study we re-analyzed EEG data of a previously published event-related potential study (Passow et al., 2014) that tested younger and older adults in a dichotic listening situation. In particular, data of 25 younger (mean age: 25.8 ± 2.7; range: 22–35 years; 12 female) and 26 older adults (mean age: 70.0 ± 4.1; range: 63–76 years; 12 female) was evaluated. Both groups showed comparable educational levels (younger adults: 13.1 ± 2.3; older adults: 12.2 ± 4.2 years of education). Prior to testing, hearing acuity and sensitivity to interaural threshold differences was assessed by means of pure-tone audiometry (cf. Passow et al., 2014).

Audiometry results were used to individually adapt auditory stimuli. Hearing thresholds > 35 dB hearing loss and interaural threshold differences >10 dB precluded participation in the study. Markers of perceptual speed (Digit Symbol Substitution Test; Wechsler, 1981) and verbal knowledge (Spot-a-Word; Lehrl, 1977) were acquired to confirm the age-typically of our sample (see Table 1). All participants were healthy, right-handed, native German speakers with normal or corrected-to-normal vision who provided written informed consent and were reimbursed for participation. The study was approved by the ethics committee of the Max Planck Institute for Human Development, Berlin, Germany.

### Experimental procedure and stimuli

We assessed selective auditory attention by instructing participants to focus attention to either the left (focus left condition; FL) or right (focus right condition, FR) ear while highly similar consonant-vowel (CV) syllable pairs were presented dichotically (i.e., simultaneously one stimulus to the left and one to the right ear). Only syllables played to the cued ear should be reported while distractor stimuli should be ignored.

To indicate their response, participants selected the target syllable from six visually displayed response options (including the target, distractor and four similar novel, i.e., not presented, syllables). As a reference, we additionally included a condition in which subjects reported which of the two presented syllables they heard most clearly without directing attention to either ear (no-focus condition, NF). The NF blocks were always completed first to avoid carry-over effects from the focused conditions (Hiscock & Stewart, 1984). After the NF blocks, the focused blocks (FL, FR) were presented in one of two counterbalancing orders (i.e., ABBABAAB or BAABABBA).

Within all attentional conditions (NF, FL and FR) the perceptual saliency of the syllables was manipulated by adjusting intensity differences between ears. In particular, either the left or right syllable′s intensity was decreased by 10 dB giving rise to three conflict conditions (Left > Right (+10); Left = Right (0); Left < Right (−10); cf. Passow et al., 2014; Tallus, Hugdahl, Alho, Medvedev, & Hämäläinen, 2007; Westerhausen et al., 2009). The target syllable′s perceptual saliency thus could match the current attentional focus (e.g., focus left and left > right (+10); i.e., low conflict; LC) or favor the distractor (e.g., focus left and left < right (−10); i.e., high conflict; HC). In the intermediate no-conflict (NC) condition, the to-be attended and distractor syllable were played out with the same intensity. Conflict conditions were randomized within and across attentional focus blocks.

Twelve CV syllable pairs consisting of syllables of voiced (/b/, /d/, /g/) or unvoiced (/p/, /t/, /k/) consonants combined with the vowel /a/ served as auditory stimuli. Each pair contained two syllables with the same voicing that were matched for onset times (cf. Hugdahl et al., 2009; Westerhausen et al., 2009). To account for aging-related hearing loss, syllable pairs were presented at an individually adjusted volume (i.e., 65 dB above participant′s hearing threshold at 500 Hz; cf. Passow et al., 2014). To determine syllable discriminability, we presented the six syllables diotically (i.e., the same syllable at the same time to the left and right ear) prior to the main task (observed mean accuracy > 90 %).

In the main dichotic listening task, each of the twelve dichotic syllable pairs was presented nine times in each of the three attentional and saliency condition, summing up to a total of 972 trials that were split in blocks of 81 trials. The inter-stimulus interval varied between 3500 and 4000 ms. Stimuli were presented using E-Prime software (v. 1.1) and insert earphones (ER 3A; Etymotic Research, Inc. Elk Grove Village, IL, USA) in an electro-magnetically shielded and sound-attenuated booth.

### Behavioral analyses

To evaluate age differences in selective attention, we calculated the auditory laterality index (LI; Marshall, Caplan, & Holmes, 1975), for each Attentional focus condition (NF, FL, FR), collapsing over conflict levels. This index expresses the amount of right relative to left ear responses (i.e., LI = (Right – Left) / (Right + Left)). The LI ranges from −100 to 100 whereby negative values indicate more left ear responses and positive values index a tendency towards selecting the right ear syllable. Younger and older adult′s laterality indexes were analyzed in a two-factorial (Age group × Attentional focus (NF, FL, FR)) repeated measures analysis of variance (ANOVA). A Greenhouse-Geisser correction (Greenhouse & Geisser, 1959) for the degrees-of-freedom was applied, whenever the assumption of homogeneity of variances across factor levels was not appropriate. For analyses regarding age differences in the interaction between perceptual saliency and attentional focus, please refer to Passow and colleagues (2012, 2014).

### Electrophysiological data recording and preprocessing

During the dichotic listening task, we continuously recorded EEG data from 61 Ag/AgCl electrodes embedded in an elastic cap that were placed according to the 10-10 system using BrainVision Recorder (BrainAmp DC amplifiers, Brain Products GmbH, Gilching, Germany; Braincap, BrainVision, respectively). An electrode above the forehead (AFz) served as ground. Three additional electrodes were placed next to each eye and below the left eye to acquire horizontal and vertical electrooculograms. Data was sampled at 1000 Hz in a bandwidth between 0.01–100 Hz and online-referenced to the right mastoid while the left mastoid was recorded as additional channel. During preparation, electrode impedances were kept <5 kΩ.

EEG data processing was performed by means of the EEGLab (Delorme & Makeig, 2004) and FieldTrip (Oostenveld, Fries, Maris, & Schoffelen, 2011) toolboxes in addition to custom-written Matlab code (The MathWorks Inc., Natick, MA, USA). For analyses, data was demeaned, re-referenced to mathematically linked mastoids, down-sampled to 500 Hz and band-pass filtered (0.2–125 Hz; fourth order Butterworth). A multi-step procedure was applied to purge data of artifacts: First, data was visually screened for periods of excessive muscle activity and subsequently independent component analysis (ICA) was used to identify and remove components related to eye, muscle and cardiac activity (e.g., Jung et al., 2000). Next, data was segmented in 5 s epochs (−1.5 s and +3.5 s with respect to stimulus onset) and submitted to a fully automatic thresholding approach for artifact rejection (cf. Nolan, Whelan, & Reilly, 2010). Excluded channels were interpolated with spherical splines (Perrin, Pernier, Bertrand, & Echallier, 1989). Finally, remaining trials were again visually screened to determine successful cleansing.

Time-varying power information for each trial and electrode was then extracted by convolution of the cleaned time domain signal with a series of Morlet wavelets with a length of seven cycles (cf. Herrmann, Grigutsch, & Busch, 2005; Werkle-Bergner, Shing, Müller, Li, & Lindenberger, 2009). Time-varying power estimates were computed for frequencies between 5–30 Hz (in steps of 1 Hz) in a time window between −1s to 1.5 s with respect to stimulus onset (time bins of 2 ms), separately for correct and incorrect trials. Incorrect trials included trials in which participants picked either the distractor or a novel response option.

### Electrophysiological data analyses

#### Within subject (first level) statistics

Within younger and older subjects, we contrasted ipsi- and contralateral EEG power by means of dependent-samples *t*-tests to isolate task-related, lateralized patterns of rhythmic neural activity (i.e., hypothesized ipsi/contralateral alpha increases/decreases, respectively; cf. Sander, Werkle-Bergner, & Lindenberger, 2012; Waldhauser, Johansson, & Hanslmayr, 2012; see Figure 1, lower part). Prior to analysis, ipsi- and contralateral activity of left and right cued (FL, FR) conditions was concatenated. To counteract an unequal distribution of FL and FR trials (i.e., less correct FL than FR trials; cf. Passow et al., 2014), we iteratively selected random subsets of the available trials using a bootstrapping procedure (n_bootstraps_ = 100 iterations; n_SelectedTrials_ = lowest trial number across conditions – 1). The mean *t*-value over the 100 bootstraps served as final first level test statistic. Ipsi vs. contra (IvsC) comparisons were computed separately for correct and incorrect trials within each conflict condition, resulting in six first level statistical maps per subject (i.e. Correct-HighConflict; Correct-NoConflict; Correct-LowConflict; Incorrect-HighConflict; Incorrect-NoConflict; Incorrect -LowConflict).

#### Within age group (second level) statistics

For analyses on the group level, first level *t*-maps were averaged within subjects over conflict conditions, resulting in a single statistical map for correct and incorrect trials, respectively (For analyses regarding Attentional focus and Conflict conditions, please see Supplemental results). Next, across subjects we contrasted IvsC *t*-maps of correct and incorrect trials to identify rhythmic neural correlates associated with correct performance in each age group (see Figure 1, middle part). In particular, we calculated non-parametric, cluster-based, random permutation tests as implemented in the Fieldtrip toolbox that effectively control the false alarm rate in case of multiple testing (Maris & Oostenveld, 2007; Oostenveld et al., 2011). In short, first a two-sided, independent samples *t*-test is calculated for each spatio-spectral-temporal (channel × frequency × time) sample. Neighboring samples with a *p*-value below 0.05 were grouped with spatially, spectrally and temporally adjacent samples to form a cluster. The sum of all *t*-values within a cluster formed the respective test-statistic. A reference distribution for the summed cluster-level *t*-values was computed via the Monte Carlo method. Specifically, in each of 10,000 repetitions, group membership was randomly assigned, a *t*-test computed and the *t*-value summed for each cluster. Observed clusters whose test-statistic exceeded the 97.5^th^ percentile for its respective reference probability distribution were considered significant.

Significant cluster results were followed up by means of two-factorial (Ipsi/Contra × Correct/Incorrect) ANOVAs within each age group to determine whether correct performance was driven either by ipsilateral synchronization, contralateral desynchronization or both. For this, power values for ipsi- and contralateral channels were extracted and averaged over significant spatio-spectral-temporal samples for correct and incorrect trials for each subject.

#### Across age group (third level) statistics

After identifying rhythmic neural correlates relating to correct performance within each age group, we investigated whether these differed reliably across groups. In other words, significant second level cluster results were tested for age-differences on a third level (see Figure 1, upper part). In particular, we contrasted the difference between correct and incorrect trials (i.e., Correct – Incorrect) in lateralized rhythmic neural activity (i.e., IvsC) across groups. Third level analyses were restricted to spatio-spectral-temporal samples that reached statistical significance on the second level. Comparisons were evaluated based on non-parametric Mann–Whitney–Wilcoxon tests (averaging over samples within participants). To investigate whether age-differences are more pronounced in pre- or post-stimulus time periods, significant third level tests were followed up by Mann–Whitney–Wilcoxon tests restricted to the respective time windows.

#### Single trial statistics

To explore the behavioral relevance of moment-to-moment fluctuations in lateralized rhythmic neural activity, mixed-effects logistic regression analyses were performed (i.e., a generalized linear mixed-effects model; GLME). We used maximum likelihood with Laplace approximation to estimate model parameters as implemented in Matlab′s statistics and machine learning toolbox.

In particular, single trial lateralized rhythmic neural activity (IvsC) in an electrode-, frequency- and time-range determined on a group level (significant second level statistics cluster for YA and OA) was used to predict each trial′s accuracy (correct/incorrect, i.e., a binomially distributed response). Only trials in which participants were cued to focus attention on one ear (i.e. FL: FR) were included in the analyses (on average 531 trials per subject; SD = 50.851 trials). By design, there is no a-priori defined correct/incorrect response for NF trials. EEG data was z-scored within subjects across trials before analyses in order to facilitate the interpretation of parameter estimates. Outlier trials with z-scores > 3 or < −3 were dropped from analyses (3.106 % of all trials). Age group and trial condition (Attentional focus (FL, FR) and Conflict (LC, NC, HC) condition) were added as additional fixed effects. To investigate age-group specific effects of lateralized rhythmic neural activity, we allowed for Age group × EEG interactions. Further, to account for overall performance differences between subjects, we included a random-effect for intercept grouped by subject (ID).

Goodness of fit of the logistic regression model was assessed by comparisons to a constant only (baseline) and fixed-effects only model. Further, to evaluate whether including rhythmic neural activity predictors significantly improved the model, we compared the full model to a restricted model without EEG predictors using a likelihood-ratio (LR) test.

## Notes

**Author note** This work was supported by Grants from the Deutsche Forschungsgemeinschaft (DFG) MWB (WE 4269/5-1), SCL (LI879/18-1), SP (PA2972/1-1). MW-B received support from Jacobs Foundation (Early Career Research Fellowship 2017-2019). Further support was provided by a grant to SCL (FZK 01GQ1424D) of the Bundesministerium für Bildung und Forschung (BMBF). MJD is a fellow of the International Max Planck Research School on the Life Course (LIFE; http://www.imprs-life.mpg.de/en) and recipient of a stipend from the Sonnenfeld-Foundation (http://www.sonnenfeld-stiftung.de/en/).

## References

Ahveninen, J., Huang, S., Belliveau, J. W., Chang, W.-T., & Hämäläinen, M. (2013). Dynamic Oscillatory Processes Governing Cued Orienting and Allocation of Auditory Attention. Journal of Cognitive Neuroscience, 25(11), 1926–1943. https://doi.org/10.1162/jocn_a_00452

Anderson, S., White-Schwoch, T., Parbery-Clark, A., & Kraus, N. (2013). A dynamic auditory-cognitive system supports speech-in-noise perception in older adults. Hearing Research, 300, 18–32. https://doi.org/10.1016Zj.heares.2013.03.006

Banerjee, S., Snyder, A. C., Molholm, S., & Foxe, J. J. (2011). Oscillatory Alpha-Band Mechanisms and the Deployment of Spatial Attention to Anticipated Auditory and Visual Target Locations: Supramodal or Sensory-Specific Control Mechanisms? Journal of Neuroscience, 31(27), 9923–9932. https://doi.org/10.1523/JNEUROSCI.4660-10.2011

Bauer, M., Kennett, S., & Driver, J. (2012). Attentional selection of location and modality in vision and touch modulates low-frequency activity in associated sensory cortices. Journal of Neurophysiology, 107(9), 2342–2351. https://doi.org/10.1152/jn.00973.2011

Bauer, M., Stenner, M.-P., Friston, K. J., & Dolan, R. J. (2014). Attentional Modulation of Alpha/Beta and Gamma Oscillations Reflect Functionally Distinct Processes. Journal of Neuroscience, 34(48), 16117–16125. https://doi.org/10.1523/JNEUROSCI.3474-13.2014

Craik, F. I. M., & Bialystok, E. (2006). Cognition through the lifespan: Mechanisms of change. Trends in Cognitive Sciences, 10(3), 131–138. https://doi.org/10.1016/j.tics.2006.01.007

Deiber, M.-P., Ibañez, V., Missonnier, P., Rodriguez, C., & Giannakopoulos, P. (2013). Age-associated modulations of cerebral oscillatory patterns related to attention control. NeuroImage, 82. https://doi.org/10.1016/j.neuroimage.2013.06.037

Delorme, A., & Makeig, S. (2004). EEGLAB: An open source toolbox for analysis of single-trial EEG dynamics including independent component analysis. Journal of Neuroscience Methods, 134(1), 9–21. https://doi.org/10.1016/j.jneumeth.2003.10.009

Engel, A. K., & Fries, P. (2010). Beta-band oscillations‒signalling the status quo? Current Opinion in Neurobiology, 20(2), 156–165. https://doi.org/10.1016/jxonb.2010.02.015

Fan, J., McCandliss, B. D., Sommer, T., Raz, A., & Posner, M. I. (2002). Testing the Efficiency and Independence of Attentional Networks. Journal of Cognitive Neuroscience, 14(3), 340–347. https://doi.org/10.1162/089892902317361886

Frey, J. N., Mainy, N., Lachaux, J.-P., Muller, N., Bertrand, O., & Weisz, N. (2014). Selective Modulation of Auditory Cortical Alpha Activity in an Audiovisual Spatial Attention Task. Journal of Neuroscience, 34(19), 6634–6639. https://doi.org/10.1523/JNEUROSCI.4813-13.2014

Frey, J. N., Ruhnau, P., & Weisz, N. (2015). Not so different after all: The same oscillatory processes support different types of attention. Brain Research, 1626, 183–197. https://doi.org/10.1016/j.brainres.2015.02.017

Getzmann, S., Golob, E. J., & Wascher, E. (2016). Focused and divided attention in a simulated cocktail-party situation: ERP evidence from younger and older adults. Neurobiology of Aging, 41, 138–149. https://doi.org/10.1016Zj.neurobiolaging.2016.02.018

Gola, M., Kamiński, J., Brzezicka, A., & Wróbel, A. (2012). Beta band oscillations as a correlate of alertness ― Changes in aging. International Journal of Psychophysiology, 85(1), 62–67. https://doi.org/10.1016/jjjpsycho.2011.09.001

Gola, M., Magnuski, M., Szumska, I., & Wróbel, A. (2013). EEG beta band activity is related to attention and attentional deficits in the visual performance of elderly subjects. International Journal of Psychophysiology, 89(3), 334–341. https://doi.org/10.1016/jjjpsycho.2013.05.007

Greenhouse, S. W., & Geisser, S. (1959). On methods in the analysis of profile data. Psychometrika, 24(2), 95–112. https://doi.org/10.1007/BF02289823

Hanslmayr, S., Staudigl, T., & Fellner, M.-C. (2012). Oscillatory power decreases and long-term memory: the information via desynchronization hypothesis. Frontiers in Human Neuroscience, 6(April), 1–12. https://doi.org/10.3389/fnhum.2012.00074

Herrmann, C. S., Grigutsch, M., & Busch, N. A. (2005). EEG oscillations and wavelet analysis. In T. C. Handy (Ed.), Event-related potentials: A methods handbook (pp. 229–259). Cambridge: MIT Press. https://doi.org/261589

Hiscock, M., & Stewart, C. (1984). The effect of asymmetrically focused attention upon subsequent ear differences in dichotic listening. Neuropsychologia, 22(3), 337–351. https://doi.org/10.1016/0028-3932(84)90080-0

Hong, X., Sun, J., Bengson, J. J., Mangun, G. R., & Tong, S. (2015). Normal aging selectively diminishes alpha lateralization in visual spatial attention. NeuroImage, 106, 353–363. https://doi.org/10.1016/j.neuroimage.2014.11.019

Hugdahl, K., Westerhausen, R., Alho, K., Medvedev, S., Laine, M., & HÄmÄläinen, H. (2009). Attention and cognitive control: Unfolding the dichotic listening story: Cognition and Neurosciences. Scandinavian Journal of Psychology, 50(1), 11–22. https://doi.org/10.1111/j.1467-9450.2008.00676.x

Jensen, O., & Mazaheri, A. (2010). Shaping Functional Architecture by Oscillatory Alpha Activity: Gating by Inhibition. Frontiers in Human Neuroscience, 4(November), 1–8. https://doi.org/10.3389/fnhum.2010.00186

Jung, T. P., Makeig, S., Humphries, C., Lee, T. W., Mckeown, M. J., Iragui, V., & Sejnowski, T. J. (2000). Removing electroencephalographic artifacts by blind source separation. Psychophysiology, 37(2), 163–178. https://doi.org/10.1017/S0048577200980259

Kerlin, J. R., Shahin, A. J., & Miller, L. M. (2010). Attentional Gain Control of Ongoing Cortical Speech Representations in a &quot;Cocktail Party&quot; Journal of Neuroscience, 30(2), 620–628. https://doi.org/10.1523/JNEUROSCI.3631-09.2010

Klimesch, W., Sauseng, P., & Hanslmayr, S. (2007). EEG alpha oscillations: The inhibition-timing hypothesis. Brain Research Reviews, 53(1), 63–88. https://doi.org/10.1016/j.brainresrev.2006.06.003

Leenders, M. P., Lozano-Soldevilla, D., Roberts, M. J., Jensen, O., & De Weerd, P. (2018). Diminished Alpha Lateralization During Working Memory but Not During Attentional Cueing in Older Adults. Cerebral Cortex, 28(1), 21–32. https://doi.org/10.1093/cercor/bhw345

Lehrl, S. (1977). Mehrfachwahl-Wortschatz-Intelligenztest MWT-B. Retrieved from https://opacplus.bsb-muenchen.de/search?id=4563875&db=100

Li, S.-C., Lindenberger, U., & Sikstrom, S. (2001). Aging cognittion: From neuromodulation to representation. Trends in Cognitive Science, 5(11), 479–486.

Lindenberger, U., Lövdén, M., Schellenbach, M., Li, S. C., & Krüger, A. (2008). Psychological principles of successful aging technologies: A mini-review. Gerontology. https://doi.org/10.1159/000116114

Lindenberger, U., & Mayr, U. (2014). Cognitive aging: Is there a dark side to environmental support? Trends in Cognitive Sciences, 18(1), 7–15. https://doi.org/10.1016/j.tics.2013.10.006

Maris, E., & Oostenveld, R. (2007). Nonparametric statistical testing of EEG-and MEG-data. Journal of Neuroscience Methods, 164(1), 177–190. https://doi.org/10.1016/j.jneumeth.2007.03.024

Marshall, J. C., Caplan, D., & Holmes, J. M. (1975). The measure of laterality. Neuropsychologia, 13(3), 315–321. https://doi.org/10.1016/0028-3932(75)90008-1

Mok, R. M., Myers, N. E., Wallis, G., & Nobre, A. C. (2016). Behavioral and Neural Markers of Flexible Attention over Working Memory in Aging. Cerebral Cortex, 26(4), 1831–1842. https://doi.org/10.1093/cercor/bhw011

Nehmer, J., Lindenberger, U., & Steinhagen-Thiessen, E. (2010). Aging and Technology-Friends, not Foes. GeroPsych, 23(2), 55–57. https://doi.org/10.1024/1662-9647/a000016

Nolan, H., Whelan, R., & Reilly, R. B. (2010). FASTER: Fully Automated Statistical Thresholding for EEG artifact Rejection. Journal of Neuroscience Methods, 192(1), 152–162. https://doi.org/10.1016/j.jneumeth.2010.07.015

Oostenveld, R., Fries, P., Maris, E., & Schoffelen, J. M. (2011). FieldTrip: Open source software for advanced analysis of MEG, EEG, and invasive electrophysiological data. Computational Intelligence and Neuroscience, 2011, 1–9. https://doi.org/10.1155/2011/156869

Passow, S., Westerhausen, R., Hugdahl, K., Wartenburger, I., Heekeren, H. R., Lindenberger, U., & Li, S. C. (2014). Electrophysiological correlates of adult age differences in attentional control of auditory processing. Cerebral Cortex, 24(1), 249–260. https://doi.org/10.1093/cercor/bhs306

Passow, S., Westerhausen, R., Wartenburger, I., Hugdahl, K., Heekeren, H. R., Lindenberger, U., & Li, S. C. (2012). Human aging compromises attentional control of auditory perception. Psychology and Aging, 27(1), 99–105. https://doi.org/10.1037/a0025667

Perrin, F., Pernier, J., Bertrand, O., & Echallier, J. F. (1989). Spherical splines for scalp potential and current density mapping. Electroencephalography and Clinical Neurophysiology, 72(2), 184–187. https://doi.org/10.1016/0013-4694(89)90180-6

Rogers, C. S., Payne, L., Maharjan, S., Wingfield, A., & Sekuler, R. (2018). Older adults show impaired modulation of attentional alpha oscillations : Evidence from dichotic listening. Psychology and Aging, 33(2), 246–258. https://doi.org/10.1037/pag0000238

Sander, M. C., Werkle-Bergner, M., & Lindenberger, U. (2012). Amplitude modulations and inter-trial phase stability of alpha-oscillations differentially reflect working memory constraints across the lifespan. NeuroImage, 59(1), 646–654. https://doi.org/10.1016/j.neuroimage.2011.06.092

Tallus, J., Hugdahl, K., Alho, K., Medvedev, S., & Hämäläinen, H. (2007). Interaural intensity difference and ear advantage in listening to dichotic consonant-vowel syllable pairs. Brain Research, 1185(1), 195–200. https://doi.org/10.1016/j.brainres.2007.09.012

Tune, S., Wöstmann, M., & Obleser, J. (2018). Probing the limits of alpha power lateralisation as a neural marker of selective attention in middle-aged and older listeners. European Journal of Neuroscience. https://doi.org/10.1111/ejn.13862

Velanova, K., Lustig, C., Jacoby, L. L., & Buckner, R. L. (2006). Evidence for Frontally Mediated Controlled Processing Differences in Older Adults. Cerebral Cortex, 17(5), 1033–1046. https://doi.org/10.1093/cercor/bhl013

Waldhauser, G. T., Johansson, M., & Hanslmayr, S. (2012). Alpha/Beta Oscillations Indicate Inhibition of Interfering Visual Memories. Journal of Neuroscience, 32(6), 1953–1961. https://doi.org/10.1523/JNEUROSCI.4201-11.2012

Wechsler, D. (1981). WAIS-R manual: Wechsler adult intelligence scale-revised. New York NY: Psychological Corporation. Retrieved from https://books.google.de/books?id=7IMiNgAACAAJ

Weisz, N., Hartmann, T., Müller, N., Lorenz, I., & Obleser, J. (2011). Alpha rhythms in audition: Cognitive and clinical perspectives. Frontiers in Psychology, 2(APR), 1–15. https://doi.org/10.3389/fpsyg.2011.00073

Werkle-Bergner, M., Grandy, T. H., Chicherio, C., Schmiedek, F., Lovden, M., & Lindenberger, U. (2014). Coordinated within-Trial Dynamics of Low-Frequency Neural Rhythms Controls Evidence Accumulation. Journal of Neuroscience, 34(25), 8519–8528. https://doi.org/10.1523/JNEUROSCI.3801-13.2014

Werkle-Bergner, M., Shing, Y. L., Müller, V., Li, S.-C., & Lindenberger, U. (2009). EEG gamma-band synchronization in visual coding from childhood to old age: evidence from evoked power and inter-trial phase locking. Clinical Neurophysiology: Official Journal of the International Federation of Clinical Neurophysiology, 120(7), 1291–1302. https://doi.org/10.1016/j.clinph.2009.04.012

Westerhausen, R., Bless, J. J., Passow, S., Kompus, K., & Hugdahl, K. (2015). Cognitive control of speech perception across the lifespan: A large-scale cross-sectional dichotic listening study. Developmental Psychology, 51(6), 806–815. https://doi.org/10.1037/dev0000014

Westerhausen, R., Bless, J., & Kompus, K. (2015). Behavioral laterality and aging: The free-recall dichotic-listening right-ear advantage increases with age. Developmental Neuropsychology, 40(5), 313–327. https://doi.org/10.1080/87565641.2015.1073291

Westerhausen, R., Moosmann, M., Alho, K., Medvedev, S., Hämäläinen, H., & Hugdahl, K. (2009). Top-down and bottom-up interaction: manipulating the dichotic listening ear advantage. Brain Research, 1250, 183–189. https://doi.org/10.1016/j.brainres.2008.10.070

Wöstmann, M., Herrmann, B., Maess, B., & Obleser, J. (2016). Spatiotemporal dynamics of auditory attention synchronize with speech. Proceedings of the National Academy of Sciences, 113(14), 3873–3878. https://doi.org/10.1073/pnas.1523357113

Wöstmann, M., Herrmann, B., Wilsch, A., & Obleser, J. (2015). Neural alpha dynamics in younger and older listeners reflect acoustic challenges and predictive benefits. The Journal of Neuroscience : The Official Journal of the Society for Neuroscience, 35(4), 1458–1467. https://doi.org/10.1523/JNEUROSCI.3250-14.2015

